# Lipidomic and Ultrastructural Characterization of Cell Envelope of *Staphylococcus aureus* Grown in the Presence of Human Serum

**DOI:** 10.1101/2020.04.08.033100

**Authors:** Kelly M. Hines, Gloria Alvarado, Xi Chen, Craig Gatto, Antje Pokorny, Francis Alonzo, Brian J. Wilkinson, Libin Xu

## Abstract

*Staphylococcus aureus* can incorporate exogenous straight-chain unsaturated and saturated fatty acids (SCUFAs and SCFAs, respectively) to replace some of the normally biosynthesized branched-chain fatty acids and SCFAs. In this study, the impact of human serum on the *S. aureus* lipidome and cell envelope structure was comprehensively characterized. When grown in the presence of 20% human serum, typical human serum lipids, such as cholesterol, sphingomyelin, phosphatidylethanolamines, and phosphatidylcholines, were present in the total lipid extracts. Mass spectrometry showed that SCUFAs were incorporated into all major *S. aureus* lipid classes, *i.e*., phosphatidylglycerols, lysyl-phosphatidylglycerols, cardiolipins, and diglucosyldiacylglycerols. Heat-killed *S. aureus* retained much fewer serum lipids and failed to incorporate SCUFAs, suggesting that association and incorporation of serum lipids with *S. aureus* requires a living or non-denatured cell. Cytoplasmic membranes isolated from lysostaphin-produced protoplasts of serum-grown cells retained serum lipids, but washing cells with Triton X-100 removed most of them. Furthermore, electron microscopy studies showed that serum-grown cells had thicker cell envelopes and associated material on the surface, which was partially removed by Triton X-100 washing. To investigate which serum lipids were preferentially hydrolyzed by *S. aureus* lipases for incorporation, we incubated individual serum lipid classes with *S. aureus* and found that cholesteryl esters (CEs) and triglycerides (TGs) are the major donors of the incorporated fatty acids. Further experiments using purified Geh lipase confirmed CEs and TGs being the substrates of this enzyme. Thus, growth in the presence of serum altered the nature of the cell surface with implications for interactions with the host.

**IMPORTANCE:** Comprehensive lipidomics of *S. aureus* grown in the presence of human serum suggests human serum lipids can associate with the cell envelope without being truly integrated into the lipid membrane. However, fatty acids-derived from human serum lipids, including unsaturated fatty acids, can be incorporated into lipid classes that can be biosynthesized by *S. aureus* itself. Cholesteryl esters and triglycerides are found to be the major source of incorporated fatty acids upon hydrolysis by lipases. These findings have significant implications for the nature of the *S. aureus* cell surface when grown *in vivo*. Changes in phospholipid and glycolipid abundances and fatty acid composition could affect membrane biophysics and function and the activity of membrane-targeting antimicrobials. Finally, the association of serum lipids with the cell envelope has implications for the physicochemical nature of the cell surface and its interaction with host defense systems.

## INTRODUCTION

*Staphylococcus aureus* is a major bacterial pathogen of great versatility capable of infecting most organs and tissues in the body. Treatment of *S. aureus* infections is challenging due to the development of resistance to multiple antibiotics. Mechanistic studies of *S. aureus* pathogenesis have been an area of active investigation for several decades, but there is still a need to understand the metabolic and structural properties of the pathogen *in vivo*, which are likely to be different from those when grown *in vitro*. In order for a pathogenic bacterium to cause an infection, it must utilize nutrients available in the infection site for replication (1). In a 1960 paper entitled “The host as a growth medium”, E.D. Garber proposed that understanding the physiology of the bacterium at the infection site was of fundamental importance (2). In recent years, several studies have reported that *ex vivo* growth of *S. aureus* in body fluids such as blood, ocular fluids, and nasal secretions, has profound impact on the characteristics of the organism and genes required for growth in these environments (3-5).

One striking example of differences between *S. aureus* cells grown in conventional artificial laboratory media versus cells grown in the presence of complex host biological materials is in the fatty acid composition of the lipids of the organism. Branched-chain fatty acids (BCFAs) and straight-chain saturated fatty acids (SCFAs) comprise the entirety of the fatty acid composition of the organism in cells grown in laboratory media (6, 7). However, it has been increasingly recognized that host fatty acids, including straight-chain unsaturated fatty acids (SCUFAs), are utilized by pathogens and incorporated directly into phospholipid molecules, thereby saving the energy and carbon costs of *de novo* fatty acid biosynthesis by the type II fatty acid synthesis (FASII) pathway (8, 9). In *S. aureus*, the fatty acids are predominantly found ester-linked in the polar lipids of the organism, with major phospholipid species being phosphatidylglycerol (PG), lysyl-phosphatidylglycerol (LysylPG), and cardiolipin (CL), and major glycolipid species being diglucosyldiacylglycerol (DGDG) and monoglucosyldiacylglycerol (MGDG) (7, 10, 11).

It is generally considered that *S. aureus* is unable to biosynthesize SCUFAs, and cells grown in the presence of serum (6), liver extract (12), and human low-density lipoprotein (LDL) and egg yolk LDL (13) have been shown to contain significant amounts of SCUFAs in their fatty acid profiles. In addition, free fatty acids are incorporated into phospholipids from medium supplemented with them (14, 15). Mass spectrometry (MS) analysis suggests that PG 33:1 is a major phospholipid when *S. aureus* is grown in the presence of LDL, which is likely made up of C18:1^Δ9^ (oleic acid) at position *sn-*1 and anteiso C15:0 at position *sn-*2 based on MS fragmentation (13, 15). The major source of lipids in human serum is from LDL particles that contain cholesterol esters, unesterified cholesterol, triglycerides, and phospholipids (16) (Fig. 1). *S. aureus* secretes at least two lipases, *S. aureus* lipase 1 (Sal1) and glycerol ester hydrolase (Geh) (17-19), that release fatty acids from lipids found in serum (6), and LDL (13). These free fatty acids are then incorporated into *S. aureus* phospholipids and glycolipids through the FakA/B and PlsXY systems (15, 20), with or without further elongation via the type II fatty acid synthesis (FASII) system (Fig. 1). The two-component fatty acid kinase system (FakA/B) produces fatty acyl-phosphate via FakA that phosphorylates fatty acids bound to FakB1 or FakB2 binding proteins, which have preferential specificities for SCFAs and SCUFAs, respectively (21). The resulting fatty acyl-phosphate is then incorporated into phospholipids via PlsXY.

**Fig. 1.**
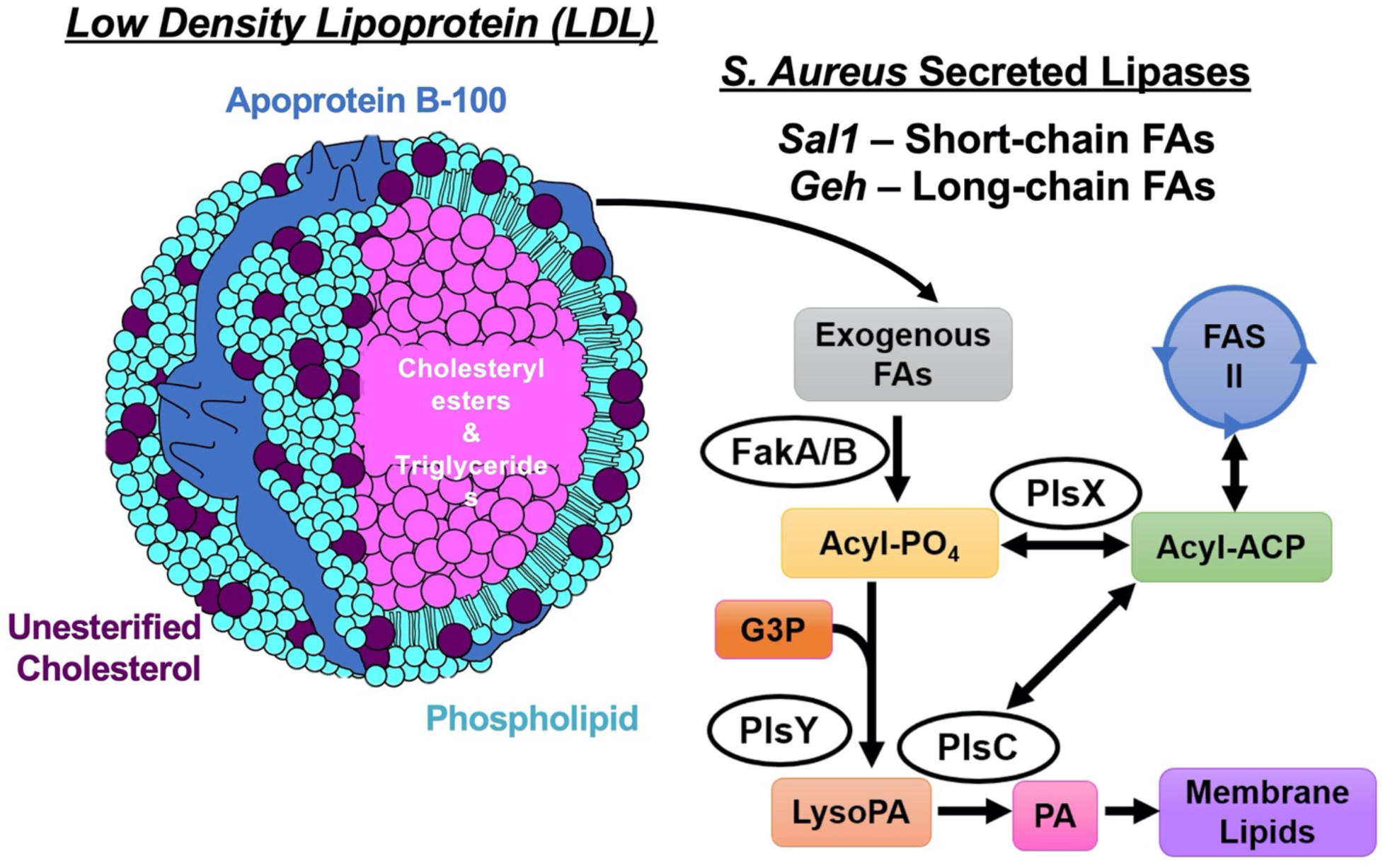
The major source of lipids in human serum is from LDL particles that contain cholesteryl esters, unesterified cholesterol, triglycerides and phospholipids. *S. aureus* secretes at least two lipases, *Sal1* and *Geh*, that release free fatty acids from lipids found in serum and LDL. These free fatty acids can be incorporated into *S. aureus* membrane lipids through the FakA/B and PlsXY systems, with or without further elongation in type II fatty acid synthesis.

However, despite previous work on utilization of exogenous fatty acids by *S. aureus*, several major questions remain. First, comprehensive lipidomic changes in the presence of exogenous lipids have not been characterized as previous studies focus on total fatty acid composition and only PGs. Second, the specific lipid classes in LDL or serum that serve as the donors of fatty acids have not been identified. Third, whether intact human serum lipids can be incorporated into the *S. aureus* membrane has not been investigated. Fourth, structural changes to the cell envelope when *S. aureus* was grown in the presence of serum have not been characterized. To answer these questions, we grew *S. aureus* in Tryptic Soy Broth (TSB) supplemented with 20% human serum and carried out comprehensive lipidomic and electron microscopic analysis of these cells. Growth of *S. aureus* in serum has the advantage of being able to mimic *in vivo* growth (22). Oogai *et al*. have shown increased expression of multiple virulence factors in *S. aureus* grown in serum (23). Supplementation of medium with blood or blood products for antimicrobial susceptibility testing of fastidious pathogens is a common practice (24, 25). The lipid composition of *S. aureus* has an impact on the interaction of the organism with the host’s defense systems (26, 27).

We demonstrated that serum-derived SCUFAs are clearly incorporated into all classes of lipids found in *S. aureus*, among which total cardiolipin levels are drastically increased when grown in the presence of serum. Interestingly, we found that serum lipids are associated with the cell envelope, which were not removed by washing with 0.9% NaCl but were removed with Triton X-100. Electron microscopy studies showed overall thickened cell envelope and loosely associated materials on the surface that were partially removable by Triton X-100. Growth in the presence of individual lipid classes indicated that cholesteryl esters and triglycerides are the major donors of the fatty acids, which is supported by studies using recombinantly expressed Geh. These findings have implications for the biological and surface properties of the organism growing *in vivo*.

## RESULTS

### S. aureus grown in serum retains serum lipids

The total extractable lipids from 1 liter of 0.9% NaCl washed cells represented about 4.6% of the dry weight of the cell, consistent with expectations (28). However, we found that total extractable lipids more than doubled (10.2%) when the cells were grown in the presence of 20% serum. Cells grown in the presence and absence of serum were subjected to comprehensive lipidomic analysis using hydrophilic interaction liquid chromatography-ion mobility-mass spectrometry (HILIC-IM-MS) (11, 29). The major lipid species observed in *S. aureus* grown in TSB included DGDGs, PGs, plasmalogen PGs (pPGs), and LysylPGs, as shown in the IM-extraction ion chromatogram (IM-XIC) in Fig. 2a. The retention time at which CLs are typically observed is noted in Fig. 2a, but CLs were below the detection limit for *S. aureus* grown in TSB (data not shown). Each class of lipids contained fully saturated fatty acids with 31 to 35 total carbons, with the species containing 33 total carbons as the most abundant species across all classes of diacyl glycerolipids (Fig. 3).

**Fig. 2.**
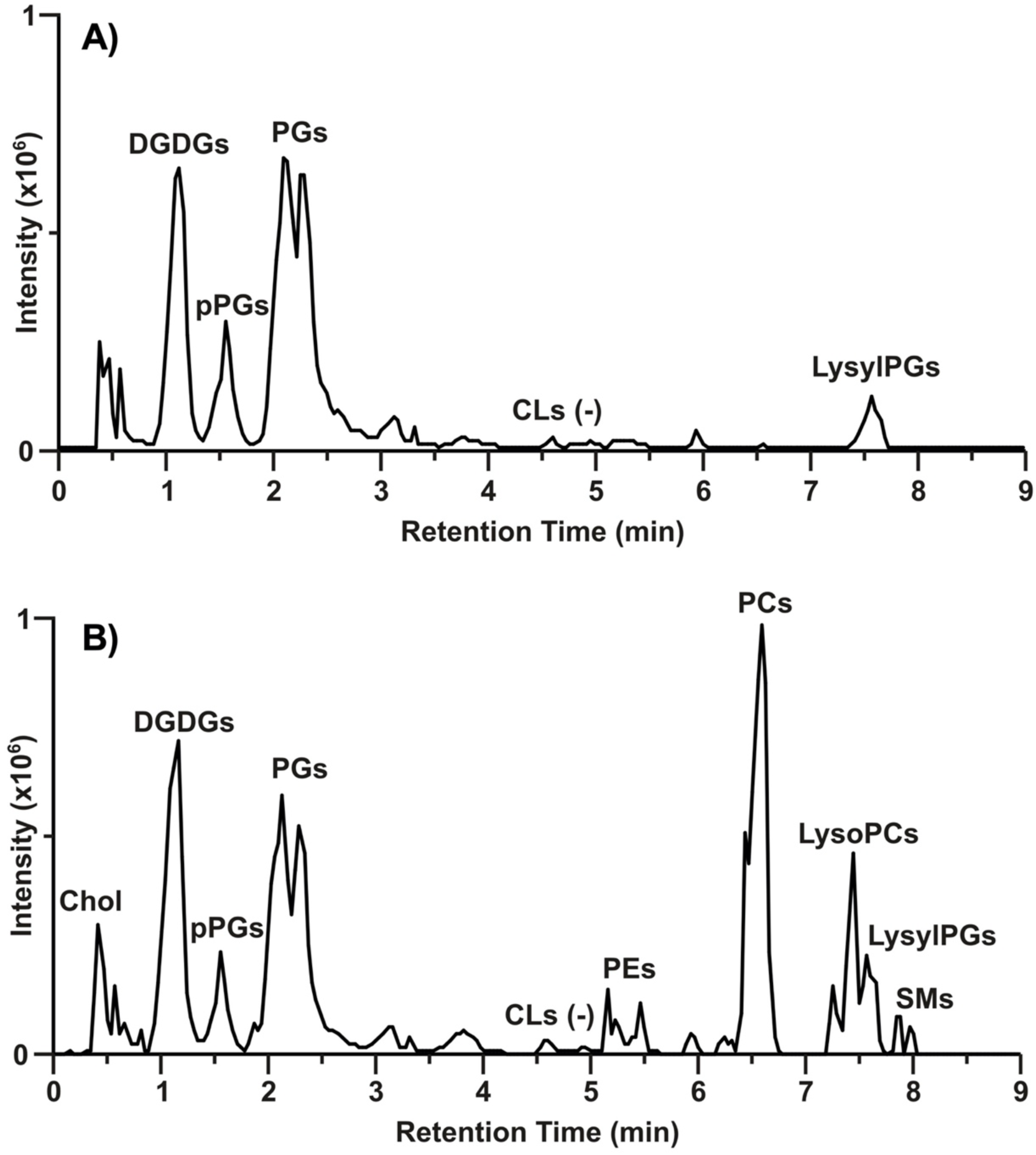
Lipid profiles of JE2 grown in A) TBS and B) TSB containing 20% human serum. Data shown are ion mobility-extracted ion chromatograms from the positive ESI analysis.

**Fig. 3.**
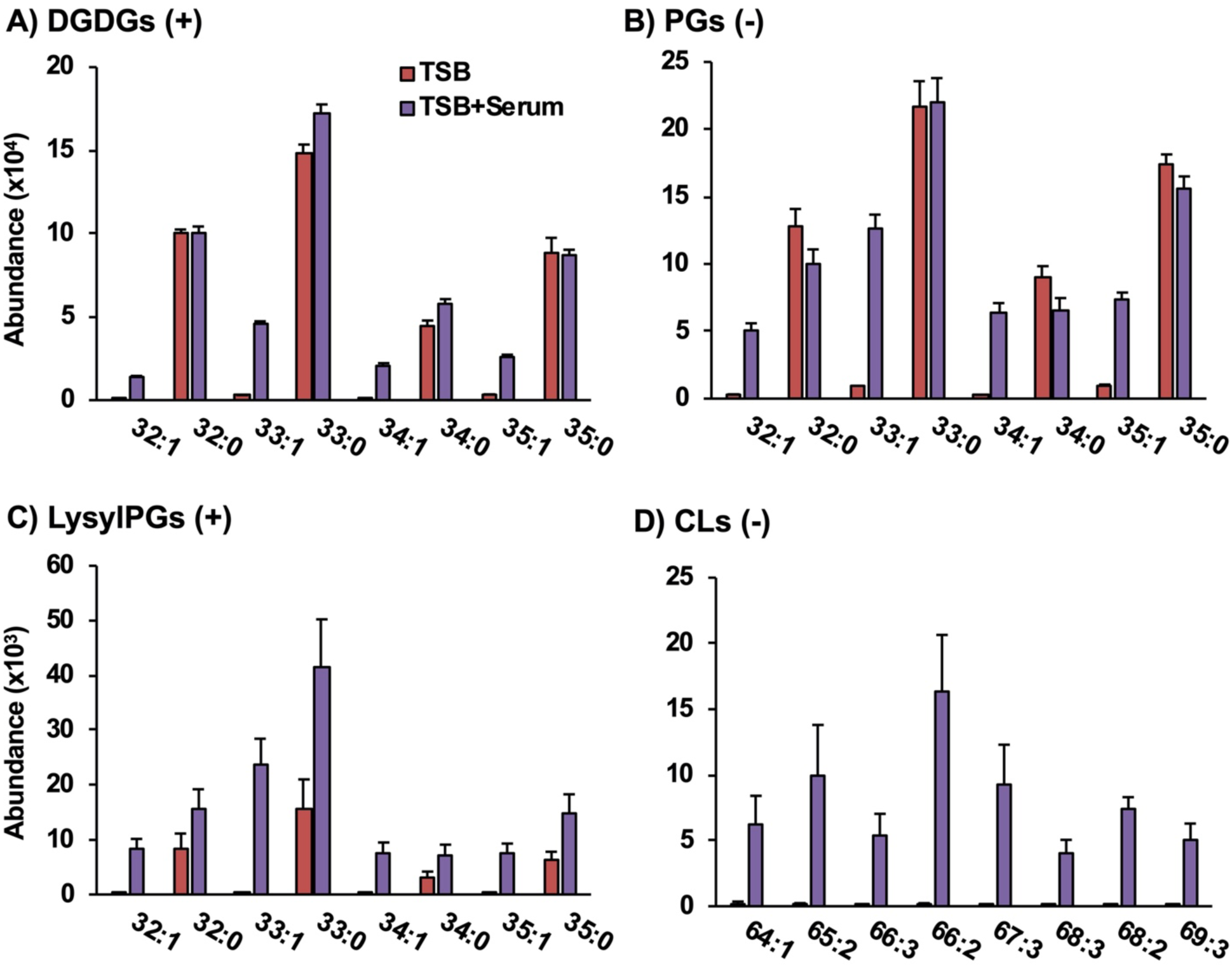
The presence of odd-carbon lipids with unsaturation are evidence that serum-derived unsaturated fatty acids are incorporated into the A) DGDG, B) PG, C) LysylPG, and D) CL lipid classes of *S. aureus.* Parentheses indicate that data is from positive (+) and negative (-) mode ESI. N = 4 per group. Statistics and detailed fatty acid composition from MS/MS experiments can be found in Supplemental Material Excel S1.

When *S. aureus* was grown in TSB supplemented with 20% human serum (TSB+Serum), the lipid profile, as shown in Fig. 2b, contained a mixture of the typical *S. aureus* lipid classes and lipids that are abundant in human serum (see Fig. S1 for lipid profile of clean TSB+Serum). The glycerophospholipids phosphatidylcholine (PC) and phosphatidylethanolamine (PE) are not produced by *S. aureus*, nor are sphingomyelins (SMs) and cholesterol (Chol). Rather, these lipids were retained by *S. aureus* from the culture through the harvesting and washing procedures.

### S. aureus grown in serum incorporates SCUFAs

The panels of Fig. 3 present the abundances of individual lipid species found in TSB only and TSB+Serum-grown *S. aureus*. Although the levels of LysylPGs (Fig. 3c) were elevated overall in the TSB+Serum condition, little to no differences were observed between TSB and TSB+Serum-grown *S. aureus* for the major fully-saturated species of DGDGs (Fig. 3a) and PGs (Fig. 3b) synthesized by *S. aureus*. However, *S. aureus* grown in TSB+Serum contained species of all major lipid classes with unsaturated fatty acids (i.e., 32:1, 33:1, 34:1, and 35:1) that were absent from *S. aureus* grown in TSB only. Specific fatty acid compositions were obtained by tandem MS (MS/MS) experiments as discussed below.

As odd-numbered carbon fatty acids are not typically observed in human serum, the occurrence of lipids with odd numbers of total carbons and one degree of unsaturation (*i.e.*, 33:1 and 35:1) strongly indicates the incorporation of a serum-derived unsaturated fatty acid into the lipids of *S. aureus*. The presence of such fatty acyl compositions in the DGDG, LysylPG, and CL species, which are not observed in human serum (see Fig. S1B), further strengthens the evidence for this incorporation.

Although CLs were not detected in *S. aureus* grown in TSB only, CLs with one to three degrees of unsaturation were present in the lipid profiles of *S. aureus* grown in TSB+Serum (Fig. 3d). The most abundant CL was CL 66:2 with 15:0 and 18:1 being the major fatty acids (see Supplemental Material Excel S1), which was consistent with the high abundance of PG 33:1 in the serum-grown *S. aureus*. These data indicate an enrichment of unsaturated CL species when *S. aureus* is grown in human serum. In contrast, no CLs were detected in the lipid profile of uninoculated TSB+Serum (see Fig. S1B).

Targeted MS/MS experiments were performed in negative ionization mode to confirm the fatty acid compositions of the lipid species presumed to contain SCUFAs based on *m/z*. An inventory of all the fatty acids observed for each lipid species in the data shown in Fig. 3, as well as those lipid species not shown in the figure, can be found in the Supplemental Material Excel S1. The most abundant fatty acyl composition across lipid species, containing 33 carbons and no double bonds, was determined to contain octadecanoic acid (C18:0) and pentadecanoic acid (C15:0). Based on the relative intensities of the two fatty acyl fragments, it is likely that 18:0 occupied the *sn-1* position on the glycerol backbone and 15:0 occupied the *sn-2* position because fatty acyl at the *sn*-2 position tends to fragment more easily (30). Using this same approach, it was confirmed that the lipids with 33:1 and 33:2 fatty acyl compositions contained 15:0 with C18:1 and C18:2 fatty acids, respectively, while 34:1 contained a major component with 16:0 and 18:1 fatty acids and a minor component with 20:1 and 14:0.

### Heat-killed *S. aureus* do not incorporate SCUFAs into their lipids

The above experiments were repeated using heat-killed *S. aureus* in order to determine whether the incorporation of SCUFAs and the retention of serum lipids were active or passive processes. Fig. 4 shows that the heat-killed *S. aureus* incubated in TSB+Serum did not contain the same levels of odd-carbon lipids with a degree of unsaturation as did live *S. aureus* incubated under the same conditions. The heat-kill reduced the levels of the endogenous lipid species as well, but to a much lesser extent. Much lower amounts of serum-derived lipids, such as cholesterol, PCs, and SMs, were observed from the heat-killed *S. aureus* compared to the live *S. aureus* when both were incubated in serum-supplemented TSB. These results indicate that SCUFA incorporation is an active process, presumably via the FakAB and PlsXY systems (20), and the retention of serum lipids also requires a living or non-denatured cell.

**Fig. 4.**
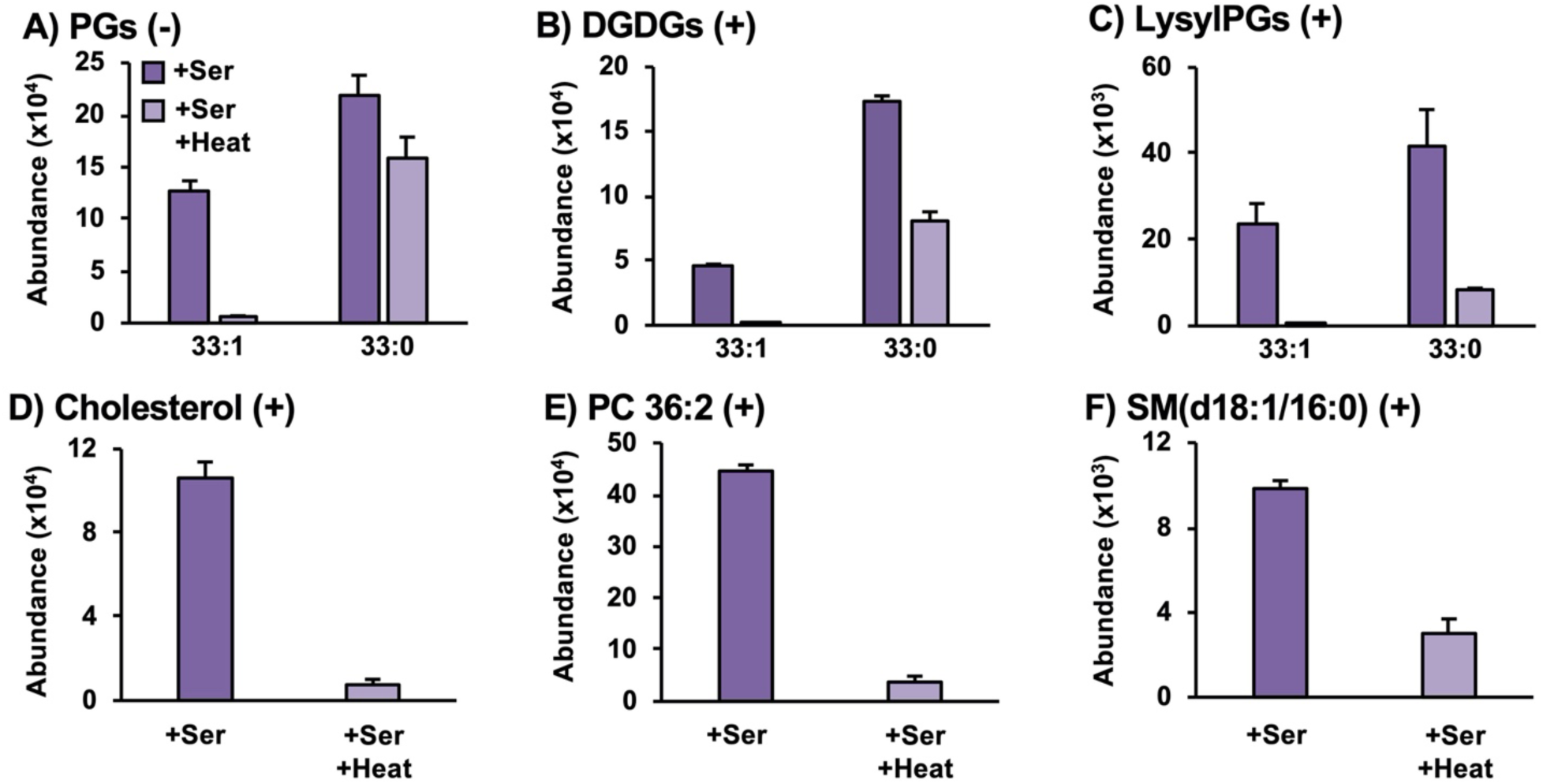
Heat-killed *S. aureus* lacks SCUFAs in A) PGs, B) DGDGs, and C) LysylPGs and retains less amounts of serum lipids such as D) cholesterol, E) PC 36:2 and F) SM(d18:1/16:0). N = 3 per group. Statistics and detailed fatty acid composition from MS/MS experiments can be found in Supplemental Material Excel S1.

### Cytoplasmic membranes isolated from TSB+Serum-grown cells retain serum lipids

Cytoplasmic membranes were isolated from *S. aureus* grown in TSB and TSB+Serum by digestion of the cell wall using lysostaphin in hypertonic sucrose followed by osmotic lysis of the protoplasts. Lipidomics was performed on washed cytoplasmic membranes. The lipid profile of the isolated membrane from TSB-grown *S. aureus* (see Fig. S2B) was consistent with the lipid profile observed for whole *S. aureus* (Fig. 2A). The cytoplasmic membrane isolated from *S. aureus* grown in the presence of serum still retained a substantial amount of serum lipids, including PCs and SMs (see Fig. S2A). The overall topography of the lipid profile was consistent with that of whole *S. aureus* cells grown in the presence of serum (Fig. 2B).

### Serum lipids are mostly removable by Triton X-100 washing

The nature of the retained serum lipids was further evaluated using a more rigorous washing procedure prior to lipid extraction. In the experiments above, pellets were washed with 0.9% NaCl solution prior to lipid extraction. To test whether the serum lipids were simply associated with the surface of the bacterium, collected *S. aureus* pellets were washed first with 0.9% NaCl, followed by a second wash with the detergent Triton X-100 (1%) to remove passively associated lipid material from the growth medium. Principle components analysis (PCA) of the resulting lipidomics data, shown in Fig. 5a, reveals that the Triton X-100 wash had a greater effect on the lipid profiles of serum-grown *S. aureus* than the *S. aureus* grown in TSB only. While PC1 clearly corresponds to the differences between TSB+serum-grown and TSB-only-grown *S. aureus*, the differences due to the NaCl and Triton X-100 washes are revealed on PC2. Along PC2, the separation between NaCl versus Triton X-100 washes for TSB+Serum-grown cells is much larger than the separation between the two washing conditions for TSB-only-grown cells. The two washing techniques had no significant effect on the abundances of the natively synthesized *S. aureus* lipids nor the incorporation of serum-derived SCUFAs into *S. aureus* lipids, as shown in Fig. 5b. However, the serum-derived PCs observed when *S. aureus* was grown in serum were nearly completely eliminated by the Triton X-100 washing (Fig. 5c).

**Fig. 5.**
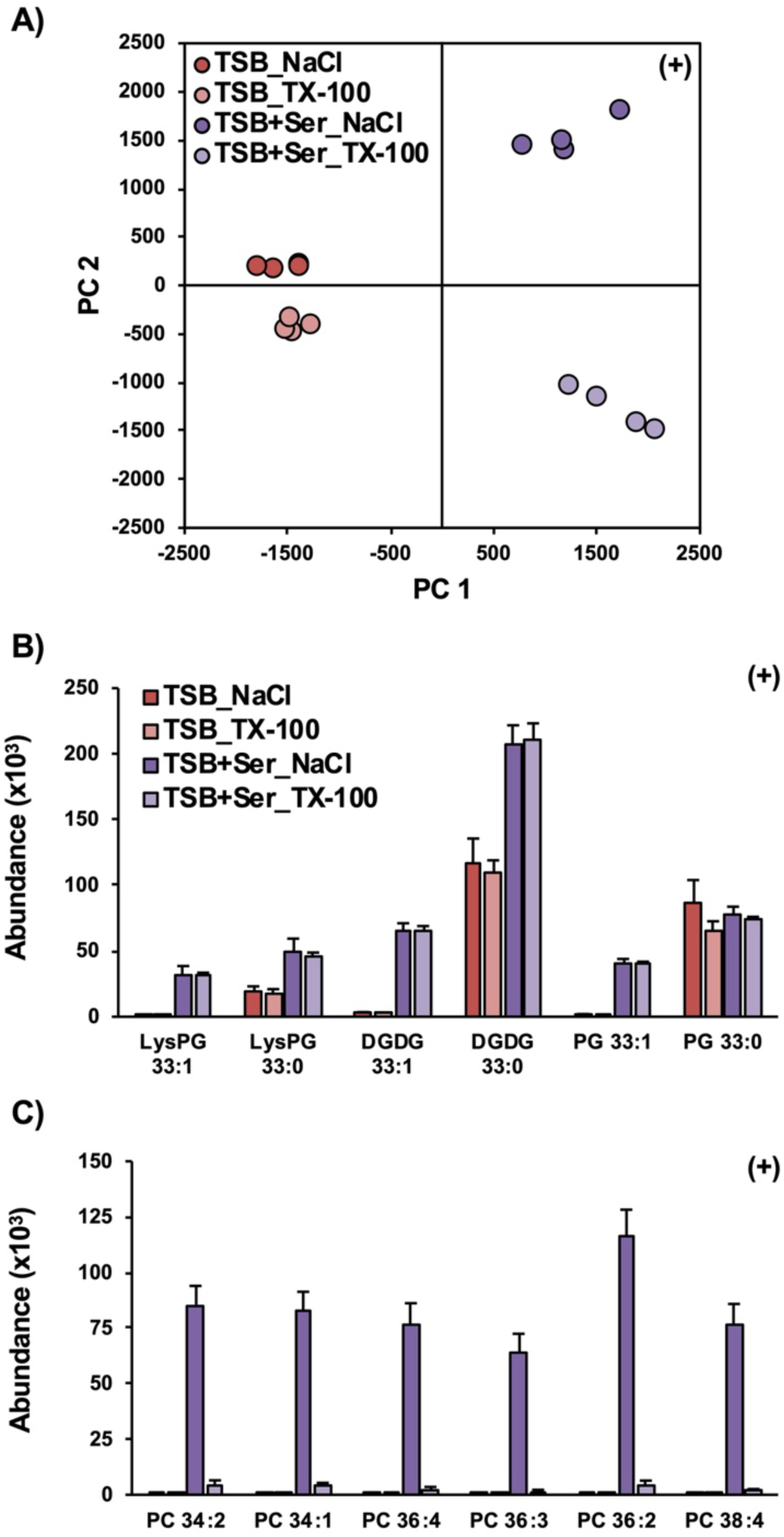
Principal component analysis of lipidomics data (A) reveals that washing pelleted *S. aureus* with TX-100 prior to lipid extraction alters the lipid profile of *S. aureus* grown in TSB supplemented with serum but not *S. aureus* grown in TSB only. B) Washing with TX-100 has no effect on the abundance of endogenous lipids or the incorporation of SCUFAs in serum-treated S. aureus. C) Pellets from serum-grown *S. aureus* treated with TX-100 prior to lipid extraction had significantly lower levels of serum lipids, such as PCs. N= 4 per group. Statistics and detailed fatty acid composition from MS/MS experiments can be found in Excel S1.

### Electron microscope studies reveal more cell clumping, associated surface material partially removable by Triton X-100, and thicker cell envelopes in serum-grown cells

Transmission and scanning electron microscopy analysis was carried out to examine the effect of serum on cell envelope structure (Fig. 6). In SEM images, *S. aureus* cells grown in the presence of serum are seen clumped together compared to cells grown in TSB only, which are more dispersed (Fig. 6A). Clumping of cells grown in TSB+Serum is consistent with observations made while handling bacterial pellets, where pellets were much harder to resuspend compared to TSB only-grown cells. Additionally, serum-grown cells display a textured cell surface unlike the smooth surface seen in TSB only-grown cells. TEM analysis revealed more detailed changes to the cell wall of serum-grown cells (Fig. 6B). TSB+Serum-grown cells appear to display a thicker cell wall and large protrusions with irregular shapes on the cell surface while TSB-grown cells again display a relative smooth cell surface. Materials at the protrusions appear to be partially removed through washing with 0.1% Triton X-100, suggesting some of these materials are associated with the cell wall. Quantitative analysis of overall cell wall thickness including the protrusion support the visual conclusions (Fig. 6C). Cell walls of TSB+Serum-grown cells are thicker than those of TSB only-grown cells regardless of washing conditions although there does not appear to be a difference between NaCl and Triton X-100 washed cells.

**Fig. 6.**
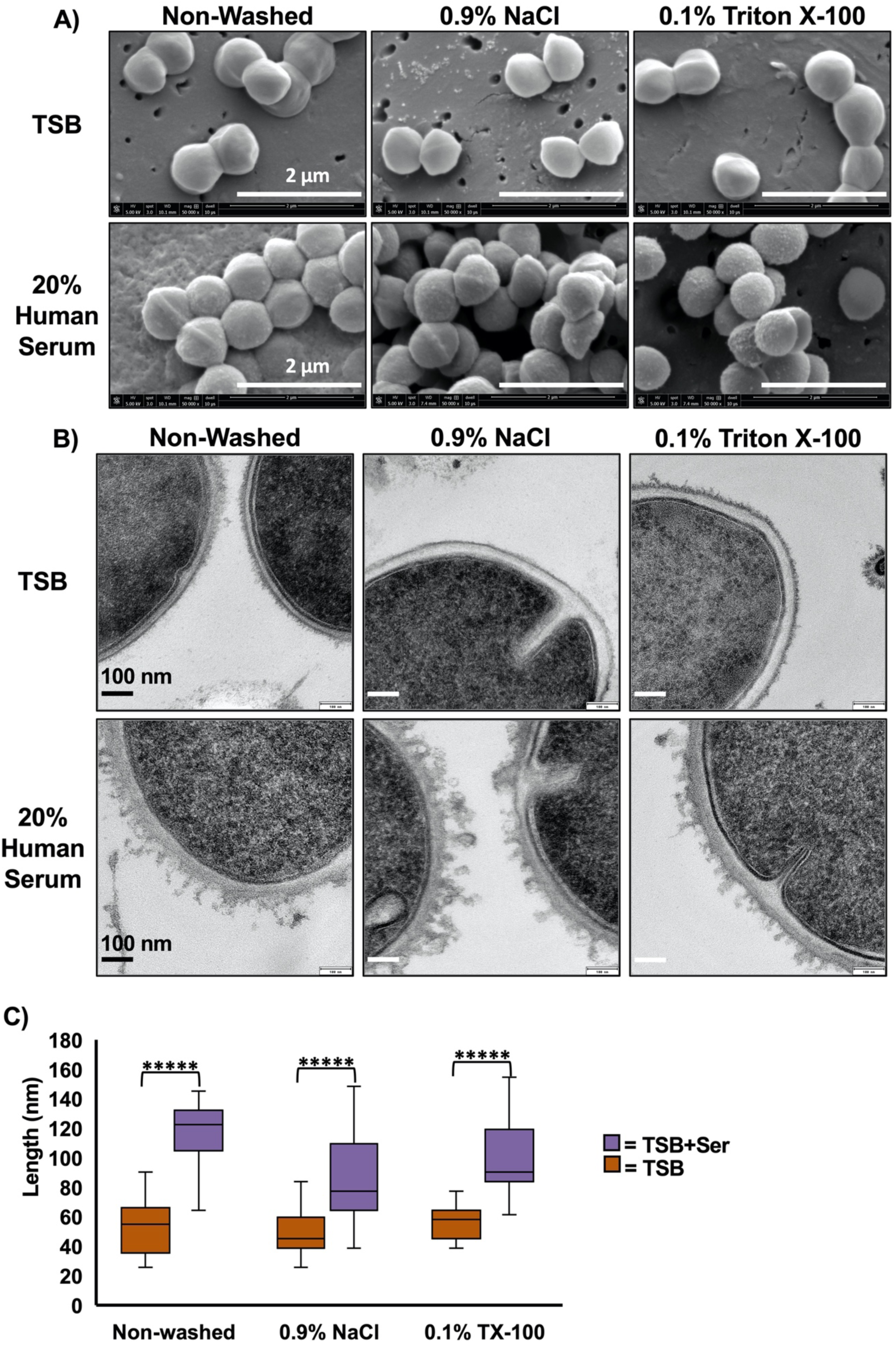
Electron microscopic analysis of *S. aureus* grown with and without human serum. (A) SEM images reveal *S. aureus* grown in the presence of human serum leads to a textured cell surface compared to the smooth cell surface of TSB-grown cells. (B) TEM images reveal that cells grown in TSB+Serum display protrusions with irregular shapes, which can be partially removed with Triton X-100 washing. (C) Quantitation of cell wall thickness reveals thicker cell walls in cells grown in TSB+Serum than those grown in TSB only, but cell wall thickness does not differ between washing with 0.9% NaCl or Triton X-100. N ≥ 21 per group. *****, *p* < 10^−6^.

### Sources of serum FAs for incorporation into S. aureus lipids

Serum is a complex mixture containing several classes of lipids that includes cholesteryl esters (CEs), triglycerides (TGs), and phospholipids (16). To evaluate which of these lipids may provide fatty acid substrates for incorporation into *S. aureus* lipids, bacteria were grown in TSB supplemented with 0.1 mM of oleic acid, cholesteryl oleate and linoleate (CEs), and extracts of PEs and PCs from chicken egg in ethanol. The lipid profiles resulting from growth of *S. aureus* with free oleic acid and the cholesteryl oleate/cholesteryl linoleate mixture were highly similar, as indicated by the tight cluster of these two sample groups in the PCA plot (Fig.7A). The ethanol treatment alone appears to increase the amount of PG extracted (Fig. 7B), but the overall effect on the lipid profile was small enough that the TSB+EtOH and TSB only samples are grouped closely in the PCA along with the egg PC-treated group. The treatment with the intact phospholipids PEs and PCs did not lead to any significant incorporation of SCUFAs into the lipids of *S. aureus*. The PE and PC extracts used in this study indeed contain significant amount of SCUFAs (PE: 18% 18:1 and 14% 18:2; PC: 32% 18:1 and 17% 18:2), suggesting they are not readily available or not good substrates of the secreted lipases under this growth condition.

**Fig. 7.**
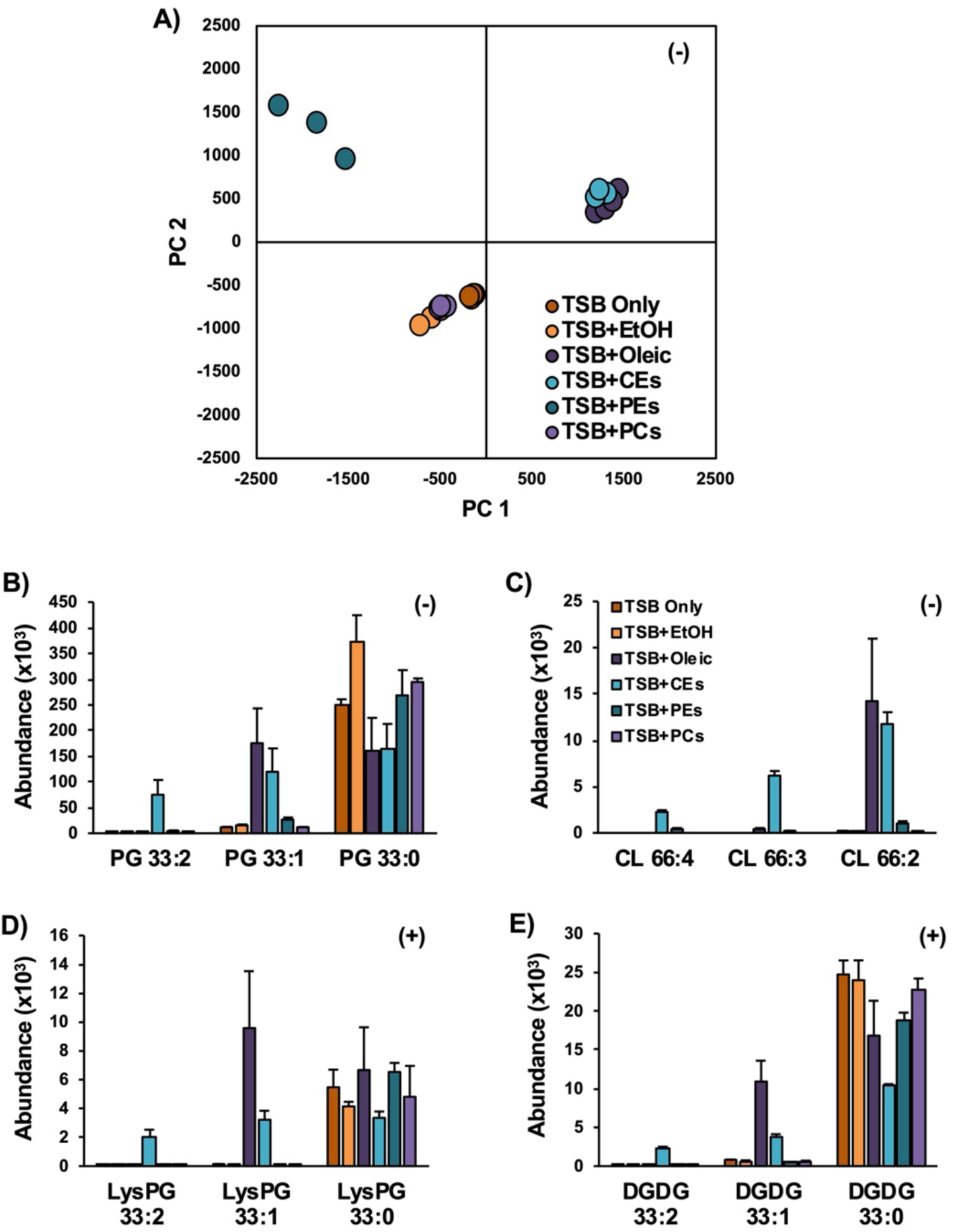
Incubations of *S. aureus* in TSB supplemented with lipid standards. A) PCA of the lipidomics data indicates that oleic acid and CEs, collectively, have similar effects on the lipid profiles of *S. aureus* relative to *S. aureus* grown in neat TSB or TSB with ethanol. The oleate and linoleate fatty acids from the oleic and CE-treated *S. aureus* are readily incorporated into B) PGs, C) CLs, D) LysylPGs, and E) DGDGs, whereas little-to-no incorporation was observed in the PE and PC-treated *S. aureus*. N = 3 per group. Statistics and detailed fatty acid composition from MS/MS experiments can be found in Excel S1.

The dramatically increased abundance of CLs with multiple degrees of unsaturation observed in the serum-grown *S. aureus* was recapitulated with the growth of *S. aureus* in TSB supplemented with oleic acid and CEs (Fig. 7C). Oleic acid and CE supplementation also resulted in the incorporation of oleate and linoleate into the major lipid classes of *S. aureus*, including PGs, LysylPG (Fig. 7D) and DGDGs (Figure 7E). Additional targeted tandem mass spectrometry was performed to confirm the fatty acid compositions of the lipid species presented in Fig. 7 as 18:2/15:0, 18:1/15:0 and 18:0/15:0, respectively (see Supplemental Material Excel S1). In a separate experiment, *S. aureus* grown in the presence of tri-oleate glyceride (TG 18:1/18:1/18:1) and tri-linoleate glyceride (TG 18:2/18:2/18:2) yielded similar results, including the high abundance of unsaturated CL species (see Fig. S3).

Evidence of *in vivo* elongation of oleic and linoleic acids into C20:1 and C20:2 fatty acids was also observed in *S. aureus* grown in TSB supplemented with oleic acid, CEs, and TGs. Figure S4 shows *S. aureus* PG, DGDG and LysylPG species with fatty acyl compositions of 35:0, 35:1 and 35:2 from *S. aureus* grown in lipid supplemented TSB. Elevated levels of 35:1 lipid species were observed from growth in the presence of oleic acid CEs and TGs. Elevated levels of 35:2 lipid species were observed in TG- and cholesteryl ester-grown *S. aureus*. Tandem MS of each lipid species individually identified the exact fatty acyl compositions for the lipids shown in Fig. S4. While the 35:0 lipid species contained 20:0 and 15:0 fatty acids, the 35:1 and 35:2 species contained 15:0 with 20:1 and 20:2 fatty acids, respectively (see Supplemental Material Excel S1). As no 20:1 and 20:2 fatty acids were supplemented into the TSB, the presence of these lipid species in *S. aureus* grown in the presence of 18:1 and 18:2 fatty acyl lipids indicated that these fatty acids were elongated prior to incorporation into diacylglycerolipids.

While oleic acid is a free fatty acid that is readily available for uptake and incorporation, the CEs and TGs contain esterified fatty acids that must undergo hydrolysis in order to generate free fatty acids. Geh is a lipase secreted by *S. aureus* with specificity for long-chain fatty acids. To evaluate the potential of Geh to generate free fatty acids, standards of CEs, TGs, PC, and PEs containing oleic or linoleic acids were incubated with purified Geh. Fig. 8 shows the abundances of free fatty acids in the supernatants following the incubation of Geh with lipid standards. As seen in the figure, despite a consistently high background level of oleic acid, CEs, PC, PE, and TG containing oleic acid yielded levels of free oleic acid higher than the background level taken from lipids that did not contain oleic acid. On the other hand, higher levels of free linoleic acid were only observed from incubation of Geh with CE, PC, and TG containing linoleic acid. The observation of PCs and PEs being substrates of Geh *in vitro*, but not donors of fatty acids *in vivo*, may be due to the different incubation conditions with the former being in 1x PBS with 10% isopropanol while the latter being in TSB with less than 1% ethanol.

**Fig. 8.**
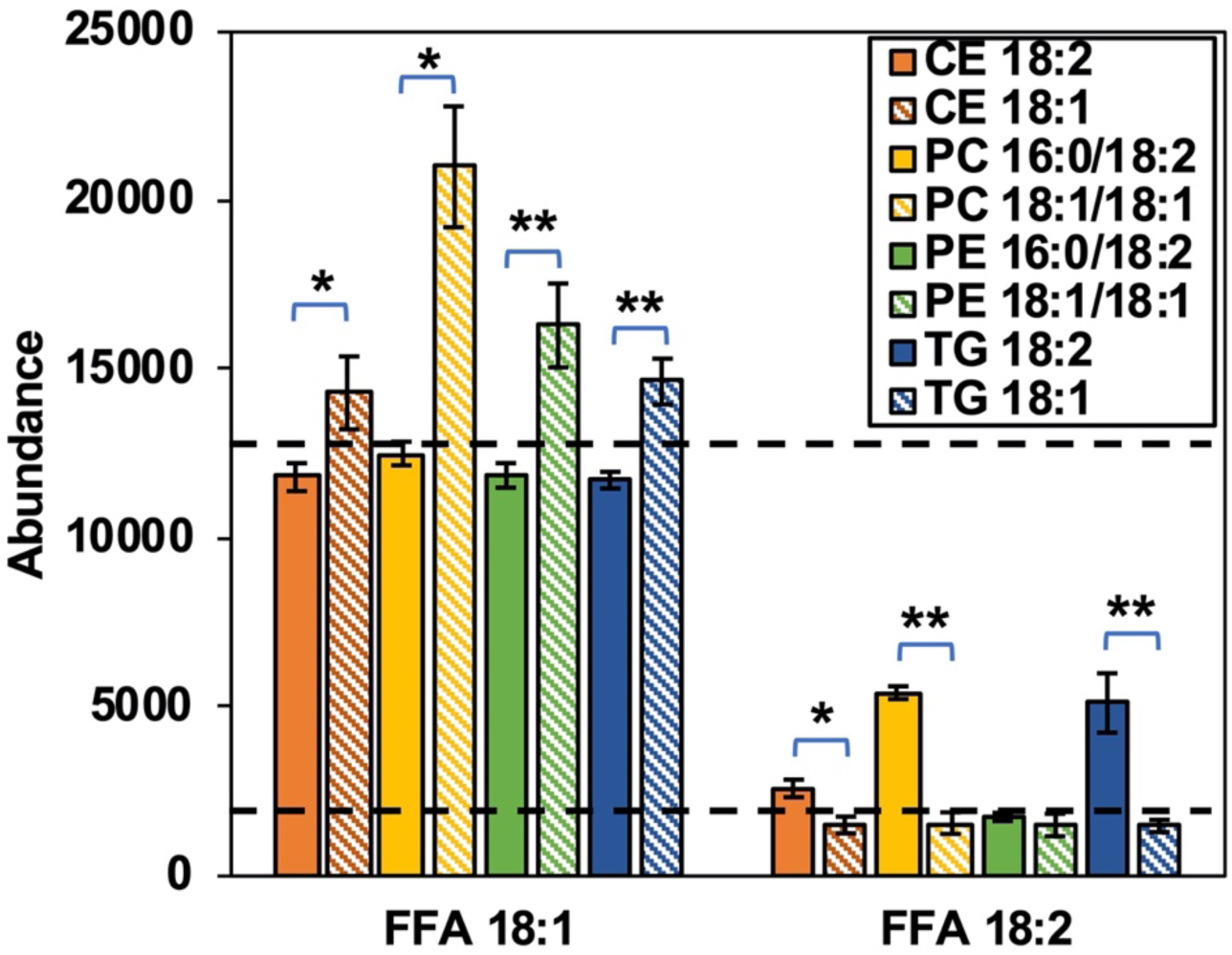
The relative abundances of free oleic (FFA 18:1) and free linoleic (FFA 18:2) resulting from the incubation of purified Geh with cholesteryl esters, phospholipids, and triglycerides containing oleic and linoleic acids. N = 3 per group. Statistics were carried out using Student’s *t*-test. *, *p* ≤ 0.05; **, *p* ≤ 0.01.

## DISCUSSION

### Increased overall lipid content in *S. aureus* grown in the presence of human serum

In 1971, Rédai et al. reported total extractable lipids comprised 20% of the dry weight of the organism grown in broth supplemented with 20% human serum (31). In this study, we found that the total extractable lipids of serum-grown *S. aureus* more than doubled (10.2% vs. 4.6%) compared to cells grown in TSB alone. This large increase in lipid content in the presence of human serum suggest host-derived lipids could be associated with or incorporated into *S. aureus* cell envelope.

### Incorporation of host fatty acids into *S. aureus* lipids

We have previously shown that SCUFAs became about 25% of the total fatty acid profile of *S. aureus* grown in 100% total bovine serum (6). Delekta et al. grew *S. aureus* in the presence of human LDL and analyzed the PG species produced under these conditions by mass spectrometry (13). PG species containing C16:1, C18:1, C18:2, and C20:1 were observed. The most abundant PG species were PG 33:1, 35:1, and 36:2. Gruss and co-workers found that addition of exogenous fatty acids promotes resistance to FASII antibiotics by *S. aureus* and selection of resistant strains that bypass FASII inhibition (12, 32). The same group showed that exogenous fatty acids could occupy both the *sn*-1 and *sn*-2 positions of PG when cells were grown in Brain Heart Infusion broth supplemented with C14:0, C16:0, and C18:1, or serum (33). This seemingly disproves the essentiality of the requirement for biosynthesized fatty acid anteiso C15:0 at the *sn*-2 position (15, 34) and undermines the viability of inhibitors of the FASII pathway as useful therapeutic agents (33). In this work, we also observed lipid species containing no C15:0, such as PG 32:1 (18:1/14:0), PG 34:1 (18:1/16:0 and 20:1/14:0), and PG 36:1 (18:1/18:0, 20:1/16:0, and 22:1/14:0), which supports the notion that anteiso C15:0 is not essential. Furthermore, we observed incorporation of SCUFA into all major classes of lipids that can be synthesized by *S. aureus* (Fig. 3, 7, and S3) and that SCUFAs can undergo elongation within *S. aureus* (Fig. S4), suggesting that host-derived fatty acids can fully participate in all fatty acid (FASII) and glycerolipid metabolic pathways.

It is particularly worth noting that the proportion of CL of the total phospholipids was drastically increased in cells grown in the presence of serum, oleic acid, and CEs (Figures 3 and 7) and these CLs contain at least one SCUFA. When grown in TSB only, no CL was detected under the same condition, including CLs with fully saturated fatty acids. CL is synthesized by condensation of two molecules of PG by CL synthase enzymes (35). The *cls*2 gene encodes the major CL synthase of the two in *S. aureus* (36, 37). Notably, all observed CLs in TSB+Serum-grown cells contain at least one SCUFA, suggesting that PGs containing a SCUFA are preferentially used as substrates of Cls2 over PGs containing fatty acids that are *de novo* synthesized by *S. aureus*.

Increased membrane CL content has been shown to be involved in decreased susceptibility to the important last-line anti-staphylococcal drug daptomycin. CL is a non-bilayer phospholipid with a small head group and four fatty acyl chains, that typically organizes in microdomains at high-curvature regions of the membrane, such as the sites of cell division and membrane fusion (9, 38-40). Daptomycin was found to attract and cluster fluid lipids in the membrane, causing membrane depolarization and delocalization of membrane proteins (41). Jiang et al. found some clinical daptomycin-resistant mutants had gain-of-function mutations in *cls*2, leading to increased CL content and decreased PG content, which then resulted in decreased daptomycin susceptibility (42). Zhang et al. have found that CL renders liposomes impermeable to daptomycin and proposed that this could be due to the prevention of flipping of the daptomycin to the inner leaflet of liposomes (43). The CL enriched membrane was also thicker than wild type membrane and resisted daptomycin lipid extraction, membrane penetration and disruption (43). In bilayer model systems, inclusion of CL has been shown to lead to increased bilayer thickness and a stiffening of the membrane, which correlates with decreases susceptibility to membrane lysis induced by helical antimicrobial peptides (44, 45). Thus, increased content of CLs in *S. aureus* grown in a host environment could result in decreased susceptibility to daptomycin and other antimicrobial peptides.

### Association of serum lipids with the cell envelope of *S. aureus*

TSB+Serum-grown *S. aureus* cells retain all major serum lipids, but these lipids are mostly removable by washing with Triton X-100. Furthermore, electron microscope images reveal that serum-grown cells have thicker cell envelopes and associated materials on their surfaces that can be partially removed by Triton X-100 washing. These observations suggest that serum lipids are associated with the cell wall, either directly as liposomes through hydrogen-bonding between the polar lipid headgroup and the cell wall or mediated by serum proteins, instead of being truly incorporated into the cell membrane. Association of serum lipids with the cell significantly decreased in heat-killed cells, suggesting the cell envelope must not be denatured for efficient association of the serum lipids. When cytoplasmic membranes were isolated from lysostaphin-induced protoplasts from serum-grown cells, the total lipid profile was very similar to that of NaCl-washed intact cells grown in TSB+Serum medium. The fact that lysostaphin-induced protoplasts, but not Triton X-100-treated cells, retain all serum lipids suggest that there is a secondary process in lysostaphin-treated cells through which the serum lipids are incorporated into the membrane.

The incorporation of serum lipids to cell wall-removed *S. aureus* is not surprising as this phenomenon has been observed in *S. aureus* L-forms. Bacterial L-forms are derived from typical bacteria, often through treatment with cell wall-active antibiotics, and lack an organized cell wall, yet they can proliferate in suitable media (46). Supplementation of medium with serum is often used to grow L-forms. Interestingly, cholesterol, cholesteryl esters, and triglycerides (all serum lipids) have been reported to be a component of the lipids of *S. aureus* L-forms although the content of PCs, PEs, and SMs was not examined (47). Nishiyama and Yamaguchi reported electron microscopic detection of complexes between the sterol-specific antibiotic filipin and cholesterol in the membrane of staphylococcal L-forms (48). Thus, the presence of cholesterol in L-forms is a precedent for our finding of this mammalian serum lipid in *S. aureus* cells and in their membranes. Interestingly, L-forms were also reported to have double the CL content of parental bacterial forms (47).

We cannot completely exclude the possibility that serum lipids, likely as small liposome vesicles, could migrate through the cell wall and directly interact with the membrane. Lee et al. show that extracellular vesicles produced from the cytoplasmic membrane of *S. aureus* can traverse the cell wall (49). Extracellular vesicles, which are delimited by a lipid bilayer and cannot replicate, are naturally released from the cells by many different organisms (50), including *S. aureus*. Coelho et al. found that the composition of extracellular vesicles from the gram-positive bacterial pathogen *Listeria monocytogenes*, grown in Brain Heart Infusion broth supplemented with 10% bovine fetal serum, were enriched in PE, sphingolipids and triacylglycerols (51). Although it is possible that serum lipids can cross the cell wall in the other direction and insert into the membrane, the fact that Triton X-100 can effectively remove these lipids make this hypothesis less likely.

### Cell surface and interaction with host defense systems

Incorporation of SCUFAs into *S. aureus* membrane has been shown to impact host-pathogen interactions. Lopez et al. showed that incorporation of *cis* SCUFAs from the host into membrane phospholipids activated the type VII secretion system for multiple virulence factors (26). On the other hand, Nguyen et al. demonstrated that SCUFAs C16:1, C18:1, and C18:2 were taken up, elongated, and incorporated into membrane phospholipids and the lipid moiety of lipoproteins. This led to an increased recognition of the *S. aureus* by the innate immune system dependent on Toll-like receptor 2 (27). However, it is also plausible the association of human serum-derived lipids with the cell envelope could change the response by the host innate immune system, *i.e*., the host material-decorated cells could allow them to escape the immune system. Detailed composition, in addition to lipids, of the associated materials and their effect on host immune system would be worth further investigation.

## MATERIALS AND METHODS

### Bacterial strain and growth conditions

The studies were carried out using *S. aureus* strain JE2 derived from strain LAC USA300, a prominent community-acquired methicillin-resistant *S. aureus* strain responsible for aggressive cutaneous and systemic infections in the USA (52). The strain was grown in Tryptic Soy Broth (TSB) (BD Difco; Franklin Lakes, NJ), at 37°C with shaking (200 rpm) in 50 mL medium in 250 mL Erlenmeyer flasks. For growth in the presence of serum, TSB was supplemented with 20% heat-treated pooled gender human serum (BioIVT; Hicksville, NY). Cultures were harvested by centrifugation (9,800 x g at 4°C for 5 min) and were washed twice by resuspension and centrifugation in cold 0.9% NaCl. For treatment with lipid standards TSB was supplemented with oleic acid (Sigma-Aldrich; St. Louis, MO), cholesteryl oleate (Sigma-Aldrich; St. Louis, MO), cholesteryl linoleate (Sigma-Aldrich; St. Louis, MO), triglycerides (NuChek Prep. Inc., Elysian, MN), phosphatidylcholine egg extract (Avanti Polar Lipids; Alabaster, AB), and phosphatidylethanolamine egg extract (Avanti Polar Lipids; Alabaster, AB) in ethanol to a final concentration of 0.1 mM in TSB. For washing experiments, cells were initially washed twice in 0.9% NaCl followed by two washes in 0.1% vol/vol Triton X-100. After Triton X-100 washing the cells were washed twice with 0.9% NaCl.

### Heat-killed cells

Cells were grown in 50 mL TSB to an OD_600_ of 1.0 and harvested and washed once in 0.9% NaCl as described above. Washed cells were resuspended in 1 mL 0.9% NaCl and incubated in a 56°C water bath for 30 mins. The heat-killed cells were then added to 50 mL of sterile TSB or TSB supplemented with 20% human serum and incubated at 37°C with shaking (200 rpm) for one hour. Cells were harvested and washed twice in 0.9% NaCl before being subjected to lipidomic analysis.

### Isolation of cytoplasmic membranes

Cytoplasmic membranes were isolated from lysostaphin-induced protoplasts as described by Wilkinson et al. (53). Briefly, washed cells were resuspended in buffered hypertonic sucrose and cell walls were digested by lysostaphin treatment. The protoplasts were recovered by centrifugation and lysed by resuspension in dilute buffer. Cytoplasmic membranes were recovered by centrifugation and were washed in water.

### Lipidomic analysis

Total lipids were extracted by the method of Bligh and Dyer (54). Lipid extracts were dried under nitrogen and dissolved in 1:1 chloroform-methanol. Small aliquots were diluted into 2:1 acetonitrile/methanol for analysis. Extracts were analyzed by hydrophilic interaction chromatography (HILIC) coupled to an ion mobility-mass spectrometer (Synapt G2-Si IM-MS; Waters Corp., Milford, MA) in positive and negative modes (11, 29). Data analysis was performed with Progenesis QI (Nonlinear Dynamics; Waters Corp., Milford, MA) and lipid abundances were normalized to bacterial dry weight (11).

### Transmission and scanning electron microscopy

Samples were prepared for transmission electron microscopy (TEM) using a modified high pressure freezing/freeze substitution (HPF/FS) method as described by Hall et al. (55). Pelleted bacteria were loaded into metal specimen carriers (2 mm diameter aluminum) coated with 1-hexadecene and frozen in an HPM 010 high pressure freezer. Freeze substitution was performed in 2% OsO_4_ (Electron Microscopy Sciences; Hatfield, PA) and 0.1% uranyl acetate (Polysciences; Warrington, PA) in 2% H_2_O and 98% acetone in an FS-8500 freeze substitution system. Samples warmed and washed as described (55). Samples were infiltrated with 1:1 Polybed812 (Polysciences) resin:acetone for 24 hours, 2:1 resin:acetone for 36 hours, 100% resin for 24 hours, and then changed to fresh resin for three days. All infiltration steps were conducted on an orbital shaker at room temperature. Samples were then submerged into embedding molds with resin and hardener and baked at 60°C for two days. 50-70 nm sections were collected using a PowerTome PC ultramicrotome with a diamond knife and collected onto carbon coated copper slot grids. Sections were imaged with a Phillips CM200 TEM. For scanning electron microscopy (SEM), samples were prepared using liquid cultures by gently passing through and embedding in a 0.2 μm filter (Nuclepore). Filter-embedded bacterial samples were fixed in 2% paraformaldehyde and 2.5% glutaraldehyde in 0.1 M sodium Cacodylate buffer at pH 7.4 for four hours on ice before being washed in buffer for 10 minutes with shaking. Samples were dehydrated in increasing ethanol concentrations three times with 10 minutes of shaking each time until reaching 100% ethanol. Samples were then dried in a Tousimis 931 critical point drier in 100% ethanol and coated with 6 nm gold-palladium. Images were collected on a FEI Quanta FEG 450 ESEM.

### Expression and purification of Geh-6xHis in *E. coli* lysY/I^q^

C-terminally 6xHis-tagged *S.aureus* Geh was expressed in *E. coli* lysY/I^q^ and purified as described previously (56).

### Incubation of lipids with Geh-6xHis

Individual lipid standards (1 mM stock in isopropanol; PCs and PEs from Avanti Polar Lipids, CEs from Sigma-Aldrich Inc., and TGs from NuChek Prep.) were added to Geh-6xHis in 50 μL 1XPBS, resulting in a 100 μM:2 μM lipid:Geh-6xHis ratio. The reaction was incubated at 37°C for 2 hours and then stopped by freezing at -80°C. Lipids were extracted and analyzed as described above.

## ACKNOWLEDGEMENTS

This grant was supported by NIH grants 1R21 AI13535 to BJW and CG, R01AI136979 to LX, and R01AI120994 to FA. FA also acknowledges support from Burroughs Wellcome Fund Investigators in the Pathogenesis of Infectious Disease Award. The authors would like to thank Ursula Reuter-Carlson, Scott Robinson, and the Microscopy Suite in The Imaging Technology Group at the University of Illinois Urbana-Champaign for assistance in TEM/SEM image collection.

## LEGENDS for Supplemental Materials

**Fig. S1.** Lipid profiles of clean A) TSB and B) TSB supplemented with 20% human serum. Data shown are IM-XICs from positive ionization mode.

**Fig. S2.** Lipid profiles of isolated cytoplasmic membranes from *S. aureus* grown in A) TSB supplemented with 20% human serum and B) TSB only. Data shown are IM-XICs from negative ionization mode.

**Fig. S3.** Incubations of *S. aureus* in TSB supplemented with tri-oleate and tri-linoleate triglycerides (TG). Oleic (18:1) and linoleic acid (18:2) were incorporated into A) PGs, B) lysylPGs, C) DGDGs, and D) cardiolipins (CLs).

**Fig. S4.** Oleic and linoleic acids derived from cholesteryl esters and triglycerides can be elongated by *S. aureus*. S. aureus lipid species PGs (A, B), DGDGs (C, D) and LysylPGs (E, F) with 35 total carbons and one or two double bonds were observed when S. aureus was grown in lipid supplemented TSB. Targeted MS/MS experiments revealed these lipids contained pentadecanoic acid and eicosenoic (20:1) or eicosadienoic (20:2) acids.

**Excel S1.** Retention time, *m/z*, collision cross section, abundance, fold-changes, statistics, and fatty acid composition obtained from MS/MS fragmentation of lipids observed in experiments related to Figs. 3, 4, 5, 7, and S3.

